# Quantifying metal ion specificity of the nickel-binding protein *Cc*NikZ-II from *Clostridium carboxidivorans* in the presence of competing metal ions

**DOI:** 10.1101/2022.10.24.513497

**Authors:** Patrick Diep, Brayden Kell, Alexander Yakunin, Andreas Hilfinger, Radhakrishnan Mahadevan

## Abstract

Many proteins bind transition metal ions as cofactors to carry out their biological functions. Despite binding affinities for divalent transition metal ions being predominantly dictated by the Irving-Williams series for wild-type proteins, *in vivo* metal ion binding specificity is ensured by intracellular mechanisms that regulate free metal ion concentrations. However, a growing area of biotechnology research considers the use of metal-binding proteins *in vitro* to purify specific metal ions from wastewater, where specificity is dictated by the protein’s metal binding affinities. A goal of metalloprotein engineering is to modulate these affinities to improve a protein’s specificity towards a particular metal; however, the quantitative relationship between the affinities and the equilibrium metal-bound protein fractions depends on the underlying binding kinetics. Here we demonstrate a high-throughput intrinsic tryptophan fluorescence quenching method to validate kinetic models in multi-metal solutions for *Cc*NikZ-II, a nickel-binding protein from *Clostridium carboxidivorans*. Using our validated models, we quantify the relationship between binding affinity and specificity in different classes of metal-binding models for *Cc*NikZ-II. We further demonstrate that principles for improving specificity through changes in binding affinity are qualitatively different depending on the competing metals, highlighting the power of mechanistic models to guide metalloprotein engineering targets.

## 1 Introduction

Many proteins rely on metal ion (herein ‘metal’) cofactors to carry out their biological function [1]. Organic molecules like proteins have a strong tendency towards forming the highest affinity complexes with Cu^II^, whereas the lowest affinity is observed for complexes with Mn^II^ and Fe^II^. More generally, metal-binding affinities of wild-type proteins are predominantly dictated by the Irving-William series: Mg^II^ < Mn^II^ < Fe^II^ < Co^II^ < Ni^II^ < Cu^II^ > Zn^II^ [2, 3]. To ensure correct metalation of proteins *in vivo* for biological function despite this hierarchy, cells have specifically evolved intracellular metal-sensing systems that control their internal metal concentrations [4, 5].

The selectivity of a protein’s metal binding site is largely determined by the electronic properties of amino acid residues in the primary coordination sphere that are spatially arranged to accommodate a metal’s preferred coordination geometry [6, 7]. The secondary coordination sphere and overall protein matrix also contribute to a protein’s metal specificity [6, 8]. The dissociation constant, *K*_D_, is a thermodynamic parameter used to quantify the binding affinity for a particular metal-protein pair. These quantities are often reported as *apparent* dissociation constants under specific pH and buffering conditions [9]. Determination of dissociation constants is often accomplished by purifying the protein [10, 11], then titrating it to saturation with the metal of interest while following the response of a fluorescence probe. The probe can either be the protein itself through intrinsic fluorescence quenching [12, 13] or an added molecular probe with fluorescence properties [9, 14].

For metals that bind the protein with high affinity, it can be difficult to reliably estimate the *K*_D_ value with direct metal titration experiments [14]. In such cases, competition experiments are used to determine the *K*_D_ instead. Competition may be achieved either by adding a molecule that competes with the protein for the metal or a second metal that competes with the metal for the protein [5, 15–21]. In the latter case, an unbound protein *P* is first saturated with metal *X*, then titrated with another metal *Y*. Under such conditions, the binding competition is described by the displacement reaction *PX* + *Y* ⇌ *PY* + *X*. Some have used this technique to study the two metals’ relative affinity described by the thermodynamic equilibrium constant for the exchange reaction, given by *K*_ex_ = *K*_D(X)_/*K*_D(Y)_ (herein ‘exchange constant’). If the dissociation constant is known for one of the metals, then the other may be estimated through empirical determination of the exchange constant. The exchange constant is also used in some studies to quantify the competition between two metals for binding a given protein [17–19].

However, the quantitative relationship between exchange constant and *in vitro* protein metalation fractions in a multi-metal solution remains unclear in general. While, methodologies have been developed to calculate fractions of a protein bound to these different metals *in vivo* [4, 5], a growing area of biotechnology research considers the use of metalloproteins *in vitro* to purify specific metals from wastewater containing several metals in large excess [22–24]. A quantitative grasp of this relationship could thus enable model-guided design of metalloproteins towards more selective biotechnologies in bio-based metal purification.

Towards this end, we present an experimental approach relying on intrinsic tryptophan fluorescence quenching to evaluate the ability of candidate models to predict a protein’s binding curves in multi-metal solutions. We used this approach to validate models of binding competition for purified *Cc*NikZ-II in two-metal solutions. In solutions of Ni^II^ and Co^II^, we found a model where both metals bind a single binding site on *Cc*NikZ-II predicted the measured binding curves adequately. However, this model did not adequately predict *Cc*NikZ-II’s binding curves in solutions of Ni^II^ and Zn^II^. In this case, we found a model including two binding sites better described the data. With the validated models, we derived predictions for the competitor-bound protein fraction at thermodynamic equilibrium for each two-metal solution. We demonstrated that the exchange constant does not uniquely determine the equilibrium competitor-bound protein fraction in our models and predict that engineered changes in the *K*_D_ values for each metal would have distinct quantitative effects on this quantity, depending on the underlying binding kinetics. Together, this work delineates a new approach for quantitatively understanding metalloprotein binding specificity in multi-metal solutions.

## 2 Methods

### 2.1 Binding model for single-metal solutions

We modelled the single-metal systems as well-mixed aqueous buffer solutions containing a metalloprotein *P* and a metal species *M*. If *M* binds a single binding site on *P* with one-to-one stoichiometry (Sect. 3.1), such a system is described by the simple equilibrium reaction

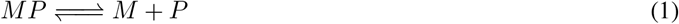

with equilibrium (dissociation) constant

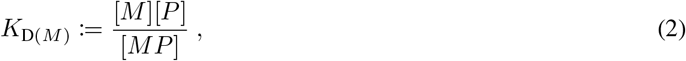

where [·] denotes an equilibrium concentration. The three equilibrium concentrations for such a system are specified by the empirical *K*_D(*M*)_ value and the mass balance equations

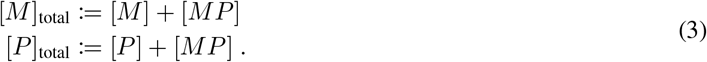

The following relationship between the measured tryptophan fluorescence quenching signal to *K*_D(*M*)_ and the (known) [*M*]_total_ and [*P*]_total_ was previously derived for this system:

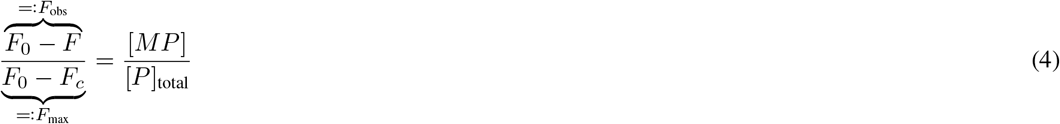

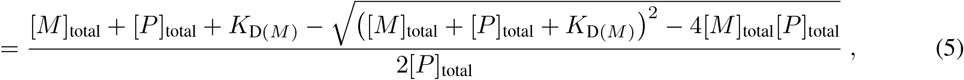

where *F*_0_ and *F_c_* are the fluorescence intensities measured when all protein molecules are unbound and bound, respectively, and *F* is the fluorescence intensity measured at occupancy fractions between 0% and 100% [25–27]. Thus, at fixed [*P*]_total_ values, experimental measurements of *F*_obs_ for varying [*M*]_total_ values can be used to estimate the parameters *K*_D(*M*)_ and *F*_max_ via curve fitting.

### 2.2 Single-site binding model for multi-metal solutions

We modelled the multi-metal systems involving a single protein binding site as well-mixed aqueous buffer solutions containing a metalloprotein *P* and *N* > 1 metal species *M_i_* (*i* = 1, …, *N*) that all bind the same site on the protein. While we restrict our experiments to two-metal systems in this work, we present our multi-metal models for arbitrarily many competing metals for generality. If each metal binds with one-to-one stoichiometry, such a system is described by the set of coupled equilibrium reactions

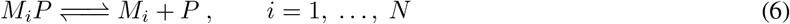

with equilibrium (dissociation) constants

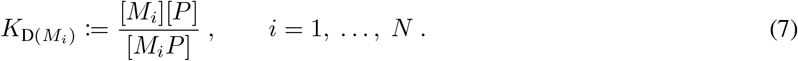

If each metal results in the same degree of fluorescence quenching upon binding, we found Eq. (4) generalized to the following (Supplementary Information Sect. S1.1):

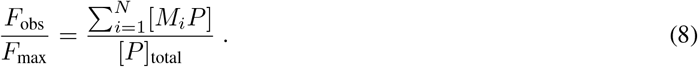

There is no general parametric solution analogous to Eq. (5) for arbitrary *N* values (Supplementary Information Sect. S1.1). However, such parametric solutions can be obtained for particular *N* ≤ 3. For larger *N*, numerical techniques have been developed to obtain numerical solutions given specified values of [*P*]_total_, *K*_D(*i*)_ and [*M_i_*]_total_ for each *i* [28, 29]. Thus, if *K*_D(*i*)_ are known from independent single-metal fluorescence quenching experiments, then Eq. (8) permits an exact theoretical prediction that may be compared against experimental data of the fluorescence signal for varied total metal concentrations at fixed total protein levels to validate the mathematical model.

### 2.3 Two-site binding model for multi-metal solutions

As in Sect. 2.2, we modelled the multi-metal systems involving a second protein binding site as well-mixed aqueous buffer solutions containing a metalloprotein *P* and *N* > 1 metal species *M_i_* (*i* = 1, …, *N*). However, here we considered the case where each metal *M_i_* to bind exactly one of two binding sites on the protein, as indicated by the sets

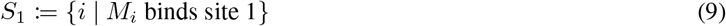

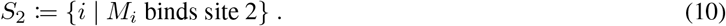

If each metal binds with one-to-one stoichiometry, such a system is described by the set of coupled equilibrium reactions

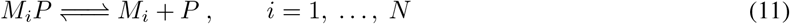

with equilibrium (dissociation) constants

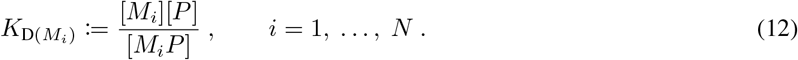

However, the presence of a second binding site permits the formation of higher-order complexes via

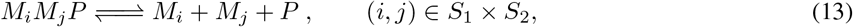

where × denotes the Cartesian product (*i.e*. 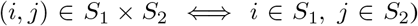. In the absence of allosteric interactions, the equilibrium constants for these reactions are constrained by

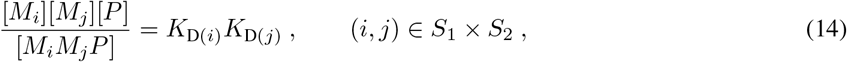

If each metal in *S*_1_ results in the same degree of fluorescence quenching upon binding, and likewise for each metal in *S*_2_ (though, not necessarily as for those in *S*_1_), we found Eq. (4) generalized to the following (Supplementary Information Sect. S1.2):

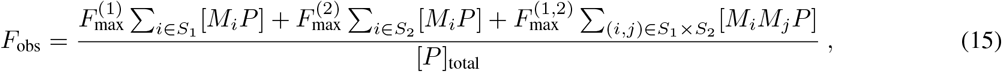

where 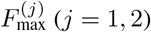 corresponds to the fluorescence signal if site *j* is fully saturated and 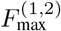 corresponds to the fluorescence signal if both site 1 and site 2 are both fully saturated. Like in the single-site model, parametric solutions can only be obtained for *N* ≤ 3, but the numerical solution algorithm in [29] may be used to obtain numerical solutions for specified values of [*P*]_total_, *K*_D(*i*)_ and [*M_i_*]_total_ for each *i* for multi-site systems. Since Eq. (15) depends on an additional parameter 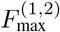 related to the formation of the second-order complexes *M_i_M_j_P*, predictions based on parameters inferred from single-metal experiments alone are not permissible. Thus, we treated 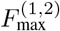 as a fit parameter in our analysis.

### 2.4 Parameter estimation and data analysis

Averages and standard deviations across three technical replicates of each fluorescence quenching experiment were calculated from the raw fluorescence data (Sect. 2.8) in `Microsoft Excel`. *K*_D_ and *F*_max_ values were then estimated from these data for single-metal experiments by least-squares optimization of Eq. (5) using the `scipy` library in in `Python 3.8.5` (Table 1). Theoretical predictions of equilibrium concentrations for the multi-metal experiments were obtained in `Wolfram Mathematica 12` by first solving for theoretical equilibrium concentrations from the mass balance and *K_D_* equations for the appropriate multi-metal model (either the single-site or the two-site multi-metal binding model, depending on the metal solution composition) and substituting the *K*_D_ values inferred from the single-metal data for each metal species (Supplementary Information Sect. S1.4). For model validation in the Ni^II^+Co^II^ systems, theoretical equilibrium concentrations were calculated for the single-site multi-metal model (Sect. 2.2) across the range of total metal and protein concentrations used in the titration and substituted into Eq. (8) with the total protein concentration and the *F*_max_ value estimated from the single-metal experiments, resulting in a predicted *F*_obs_ signal for the given experimental parameters. An analogous procedure was followed to evaluate the agreement of the single-site multi-metal model (Sect. 2.2) with the data for the Ni^II^+Zn^II^ systems. Given inadequate agreement of the single-site model with the Ni^II^+Zn^II^ data, theoretical equilibrium concentrations were also calculated for the two-site multi-metal model (Sect. 2.3) across the range of total metal concentrations used in the titration and substituted into Eq. (15) with the total protein concentration. Additionally, 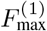 and 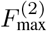 were set to the *F*_max_ values estimated from the single-metal experiments for Ni^II^ and Zn^II^, respectively. Since 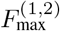 cannot be inferred from single-metal experiments, this parameter was estimated by least-squares optimization of Eq. (15) using the scipy in Python 3.8.5 (Table 2). Reduced 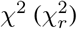 values were calculated to assess the agreement of both fitted and predicted curves with the data [30] (Table 1, Table 2, Supplementary Tables S1-S4).

**Table 1:**
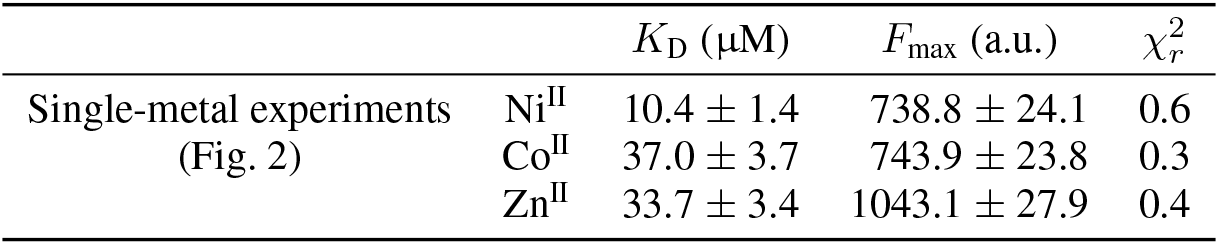
Fitted Parameters and 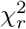 from Single-metal Experiments

**Table 2:** Fitted Parameter and 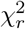 for Ni^II^+Zn^II^ Experiments

| | $F_{\max}^{(1,2)}$ (a.u.) | $\chi_r^2$ |
| --- | --- | --- |
| Ni<sup>II</sup>+Zn<sup>II</sup> experiments<br>(Fig. 4) | $1048.2 \pm 24.6$ | 3.8 |

### 2.5 DNA synthesis, cloning, and strains

The open-reading frame encoding *Cc*NikZ-II from C. *carboxidivorans* P7T (Ccar_RS11695, NCBI ID: NZ_CP011803.1) was synthesized (Twist Bioscience, San Francisco, USA) without its native signal peptide identified using SignalP 5.0 with the Gram-positive setting [31]. This was cloned into p15TV-L (AddGene ID: 26093) under the T7 promoter in-frame with the N-terminal 6xHisTag, then transformed into a LOBSTR (Kerafast, #EC1001) *Escherichia coli* strain and subsequently plated onto LB-Agar (carbenicillin, 100 μg/mL). Plasmids were sequence-verified at the ACGT Sequencing Facility (Toronto, Canada). Glycerol stocks were stored at −80 °C.

### 2.6 Protein expression and purification

This protocol is based on prior work [32]. Starter cultures of *Cc*NikZ-II were grown from glycerol stock in LB with carbenicillin (100 μg/mL) for 16 h overnight at 37 ° C with shaking. Expression cultures were started by pre-warming TB media with 5% glycerol and carbenicillin (100 μg/mL) to 37 °C before 1% v/v inoculation with the starter culture, then grown for 5 hr at 37 °C until addition of IPTG (BioShop, #IPT002) to 0.4 mM for induction. The expression cultures were then transferred to 16 °C and grown for 16 hr overnight with shaking, then pelleted with centrifugation and transferred to conical vials for one freeze-thaw cycle at −20 °C.

Frozen cell pellets were thawed and re-suspended in binding buffer (10 mM HEPES, 500 mM NaCl, 5 mM imidazole, pH 7.2) to a final volume of 100 mL, followed by addition of 0.25 g lysozyme (BioShop, #LYS702). Cell pellet mixtures were sonicated for 15 min (Q700 Sonicator, Qsonica) and clarified by centrifugation. The soluble layer (supernatant) was applied to a cobalt-charged (Co-)NTA resin (Thermo Fisher, #88221) pre-equilibrated with binding buffer in a gravity-column set-up. Bound proteins were cleansed with wash buffer (10 mM HEPES, 500 mM NaCl, 25 mM imidazole, pH 7.2) and the bound protein was collected with elution buffer (10 mM HEPES, 500 mM NaCl, 250 mM imidazole, pH 7.2). Protein concentrations were determined by Bicinchoninic acid assay (BCA), and protein purity was determined by SDS-PAGE analysis and densitometry on Image Lab 6.0 (Bio-Rad) [33, 34].

Eluted protein was combined with 0.4 mg of in-house purified TEV protease and DTT to 1 mM (BioShop, #TCE101), then transferred to a 10 kDa molecular weight cut-off dialysis bag (Thermo Fisher, #68100) for dialysis in 4 L dialysis buffer (10 mM HEPES, 1 mM TCEP, 1 g/L Chelex 100, pH 7.2) at 4 °C with gentle stirring for 24 hr. Dialyzed samples were then applied to a cobalt-charged NTA resin twice and transferred to a 10 kDa molecular weight cut-off dialysis bag for dialysis in 4 L dialysis buffer (10 mM HEPES, pH 7.2) at 4 °C with gentle stirring for 24 hr and transferred to fresh dialysis buffer for another 24 hr. Finally, samples were flash-frozen in 200 μL aliquots in liquid nitrogen before storage at −80 °C.

### 2.7 Preparation of activity buffers and metal solutions

Ni^II^ and Co^II^ solutions were prepared from hexahydrate chloride salts NiCl_2_ (BioShop, #NIC999.500) and CoCl_2_ (BioShop, #COB001.100) in non-buffered deionized MilliQ water. The concentrations were then determined by inductively coupled plasma mass spectrometry (ICP-MS) to be 73.24 mM Ni^II^ and 78.34 mM Co^II^, respectively (see Section 2.10). Ni^II^, Co^II^, and Ni^II^+Co^II^ working solutions were then made for experiments from these stock solutions using activity buffer (10 mM HEPES, pH 7.2) previously prepared from deionized MilliQ water adjusted to the desired pH using NaOH, then filter sterilized by syringe using a 0.2 μm PTFE membrane disc (PALL, #4187). The pH of the working solutions were unchanged based on visual inspection by litmus paper, which was expected based on `OLI Studio 9.6` simulations. Additionally, more than 99% of the metals were in the aqueous phase based on OLI calculations for all metal concentrations used (between 0 to 0.7 mM).

### 2.8 Generating binding curves by measuring intrinsic tryptophan fluorescence quenching

The microplate-based intrinsic tryptophan fluorescence quenching ligand-titration assay is based on prior work [32]. Binding assays were performed in black, opaque 96-well microplates (Greiner Bio-One, #655076) using the Infinite® 200 PRO (Tecan) plate reader with the following settings: fluorescence top-read, 25 °C, λ*_ex_* = 280 nm, λ*_em_* = 380 nm, and manual gain of 100.

The general procedure used to obtain binding curves first required 150 μL of 0.4 μM *Cc*NikZ-II in activity buffer to be added to a well and equilibrated for 20 min with shaking (orbital, 3mm) to 25 °C. The baseline fluorescence was monitored to ensure equilibration, from which one endpoint reading was made as the blank (*F*_0_). Step-wise titrations of metal working solution were added until saturation. At each titration step, a small aliquot (1 to 20 μL) is added by multi-channel pipette, then equilibrated for 3 min with shaking (orbital, 3mm). For larger aliquots (10 to 20 μL), the samples were equilibrated for at least 5 min with shaking (orbital, 3mm) or until the fluorescence stabilized. At each titration step, after the equilibration period, the fluorescence was measured (F). The protein and metal concentrations differ for each experiment and are described in Sects. 3.1 and 3.2. Experiments were performed in triplicates.

Mixing by pipette was avoided to prevent the introduction of air bubbles or the unintentional removal of protein due to droplets sticking to the inside of tips. To account for possible dilution effects, we included a triplicate negative control where the protein was titrated with only activity buffer using the same volumes added at each titration step. The inner-filter effect was not observed for the metal working solutions.

### 2.9 Determination of metal-protein stoichiometry by membrane filtration

A 1:100 protein-metal solution (1 mL) containing 1.7 μM *Cc*NikZ-II (0.1 μg/mL) and 170 μM Ni^II^ in activity buffer was prepared and transferred to the upper portion of a 10 kDa molecular weight cut-off concentrator (Cytiva, #28932296). This was then placed into a temperature-controlled centrifuge where it was first equilibrated for 30 min at 25 °C without disturbance, then suddenly centrifuged at 2000 rpm for 15 min. The initial permeate (labelled FT for flow-through) was transferred to a pre-weighed microfuge tube. To wash the retentate, 1 mL of activity buffer was added to the retentate and quickly centrifuged again, as described above. The first wash permeate was transferred to another pre-weighed microfuge tube. This wash process was repeated twice for a total of three washes to create three wash permeates (labelled W1, W2, and W3). The masses of FT, W1, W2, and W3 were determined by subtracting the pre-recorded initial tube weights. The volume of each fraction was calculated assuming a density of 1 g/mL. Negative controls were included where either *Cc*NikZ-II, Ni^II^, or both were absent. The experiment was performed in duplicates. The metal concentration was determined for each fraction by ICP-MS (see Section 2.10), and the protein concentration was determined by BCA assay [33], following the manufacturer protocol (Thermo Fisher, #23227).

### 2.10 Metal analysis by ICP-MS

Metal concentrations were determined by ICP-MS using a Thermo Scientific™ iCAP Q ICP-MS system in kinetic energy discrimination mode. Samples were prepared by dilution into a 2% HNO_3_ solution made from deionized MilliQ water and ultrapure HNO_3_ Optima™ (Fisher Chemical, #S020101TFIF01). Ni^II^ and Co^II^ calibration curves were created using ICP-grade Ni^II^ (Millipore Sigma, #28944) and Co^II^ (Millipore Sigma, #30329) standards. 0, 20, 100, 500, and 1000 ppb metal standards were prepared with glass volumetric flasks and pipettes previously triple-washed with 10% HNO_3_ TraceMetal™ (Fisher Chemical, #S010101TFIQ03).

## 3 Results & Discussion

### 3.1 Single-metal experiments to determine stoichiometry and *K*_D_ of *Cc*NikZ-II

*Cc*NikZ-II is homologous to the solute-binding component of *Campylobacter jejuni*’s nickel-specific ABC transporter, which is involved in enzymatic processes requiring nickel-based cofactors [35, 36]. Indeed, in prior work we demonstrated *Cc*NikZ-II binds to Ni^II^, as expected [32]. Additionally, our prior work suggests Ni^II^ binds *Cc*NikZ-II with 1:1 stoichiometry based on a fitted Hill coefficient [32]. Here, we sought to directly measure the ratio of protein-bound Ni^II^ to Ni^II^-bound protein to verify this inference. *Cc*NikZ-II from *C. carboxidivorans* was recombinantly expressed in *E. coli* cells and affinity purified to over 95% homogeneity (Sects. 2.5, 2.6). To determine the Ni^II^ binding stoichiometry of *Cc*NikZ-II, the purified protein was pre-incubated in excess Ni^II^ (1:100), and the unbound ions were washed out using centrifugal filters with a 10 kDa molecular weight cut-off (Sect 2.9). Virtually no Ni^II^ or *Cc*NikZ-II protein was observed in controls where they were not added (Fig. 1AB, Supplementary Table S3). The empirical stoichiometric ratio between Ni^II^ and *Cc*NikZ-II across the two technical replicates was 1.18 and 0.91 (Fig. 1C), which provided experimental evidence to support the modelling assumption of 1:1 metal-binding to *Cc*NikZ-II.

**Figure 1:**
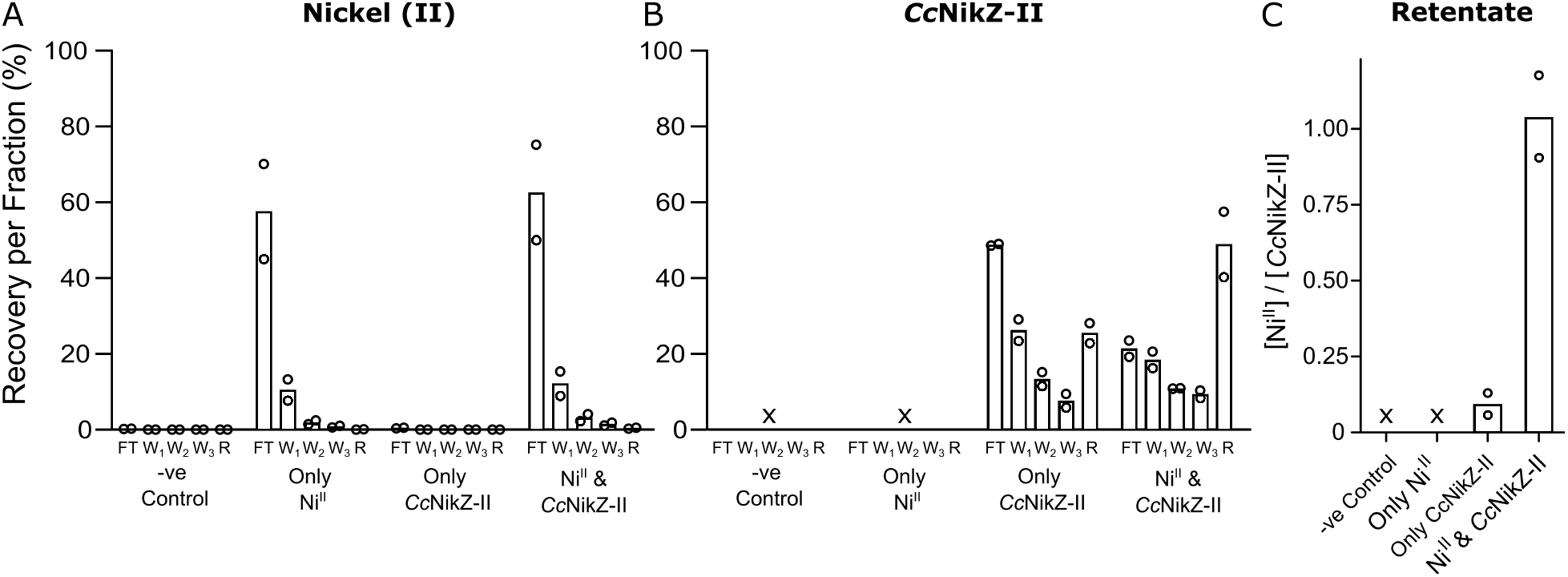
Determination of metal stoichiometry using membrane filtration. FT - Flow through, W1 - Wash 1, W2 - Wash 2, W3 - Wash 3, R - Final Retentate. Ni^II^ recovery percentages for each group were calculated relative to the initial Ni^II^ concentration in the Only Ni^II^ group. Similarly, *Cc*NikZ-II recovery percentages for each group were calculated relative to the initial *Cc*NikZ-II concentration in the Only *Cc*NikZ-II group. “x” denotes where calculations could not be made due to the protein concentration being below the limit of detection. (A) Recovery of Ni^II^ per fraction determined by ICP-MS, (B) Recovery of protein per fraction determined by BCA assay, (C) Final molar stoichiometry of Ni^II^ to *Cc*NikZ-II in retentate after W3. Experiments were performed in duplicates. Bars indicate the average across the replicates, open circles correspond to the raw data.

We then determined the *K*_D_ values of *Cc*NikZ-II for each of Ni^II^, Co^II^, and Zn^II^ using the microplate-based intrinsic tryptophan fluorescence quenching ligand-titration assay and fitting the data to Eq. (5) (Fig. 2). Similar to results in Diep *et al*. (2020) [32], we observed *Cc*NikZ-II had higher affinity for Ni^II^ than Co^II^ or Zn^II^ (Table 1), which is consistent with the Irving-William series (see Sect. 1). We also note that *Cc*NikZ-II had a similar *F*_max_ value for both Ni^II^ and Co^II^, which suggested the polar environments surrounding *Cc*NikZ-II’s tryptophan moieties undergo similar changes upon binding to either metal, consistent with a single-site binding model (see Sect. 3.2). Conversely, *Cc*NikZ-II’s *F*_max_ value differed significantly between Ni^II^ and Zn^II^ experiments, which was previously observed in [32].

**Figure 2:**
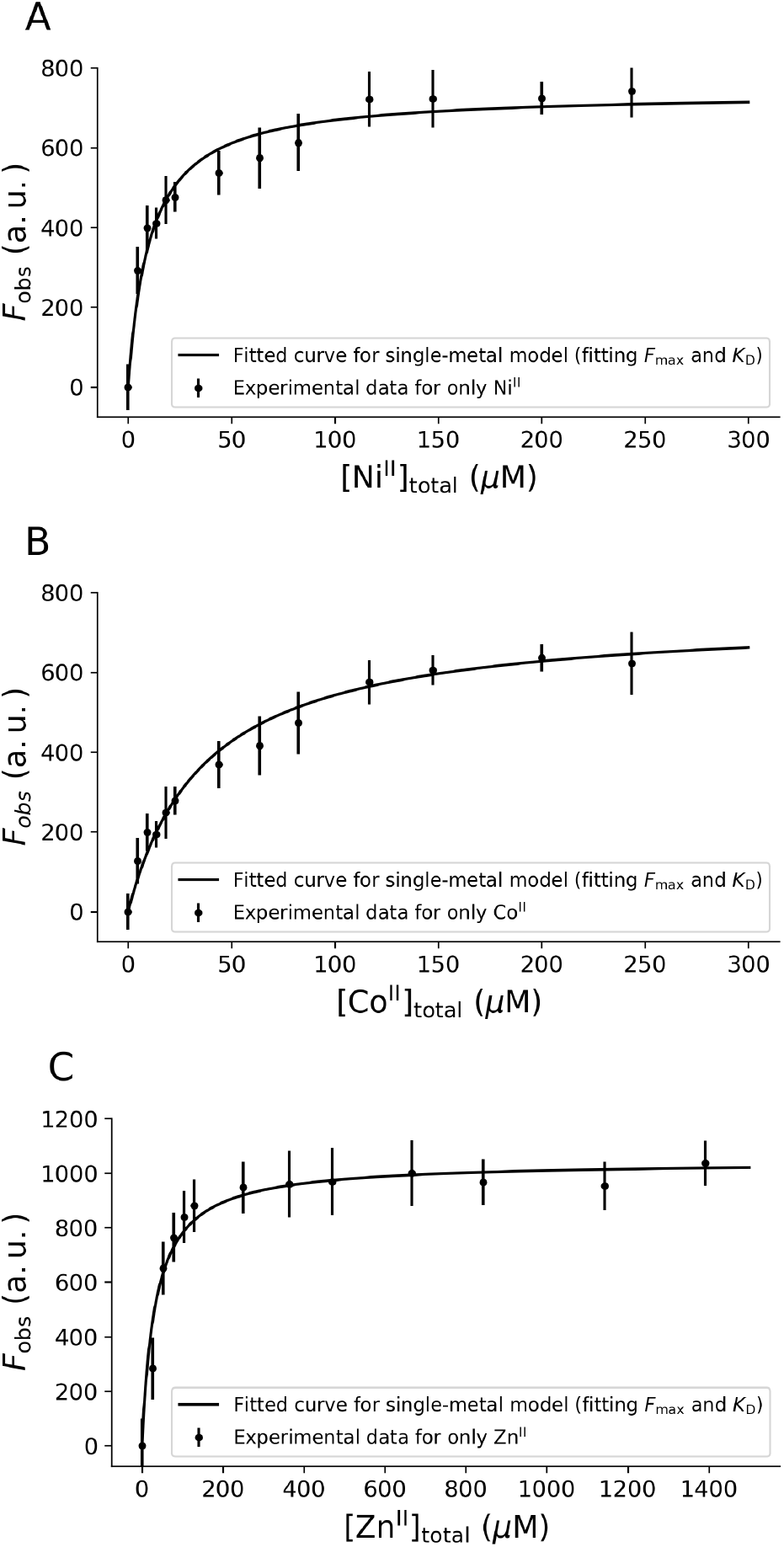
Independent binding curves of *Cc*NikZ-II separately titrated with Ni^II^ (A), Co^II^ (B), or Zn^II^ (C). The binding curves shown correspond to Eq. (5) evaluated for the *K*_D_ and *F*_max_ inferred via least squares optimization to the data (Table 1). Experiments were performed in triplicate. The dots and error bars denote the mean and standard deviation, respectively, across the technical replicates for each titration step.

### 3.2 Model validation by comparison of predicted binding curves to data in two-metal solutions

After estimating the *K*_D_ and *F*_max_ values for each of Ni^II^, Co^II^, and Zn^II^, we sought to assess the ability of our models to predict binding curves for *Cc*NikZ-II in the presence of Ni^II^ and Co^II^ or Ni^II^ and Zn^II^ at varied total concentrations of the competitor (Co^II^ or Zn^II^) relative to the total Ni^II^ concentration.

First, we assessed the ability of the single-site model (Sect. 2.2) to predict binding curves for *Cc*NikZ-II in solutions with Ni^II^ and Co^II^ (Fig. 3). To obtain such binding curves, we titrated the protein with metal stock solutions containing a mixture of the two metals at total metal ratios *f*_Co_ := [Co^II^]_total_/[Ni^II^]_total_ = 0.1, 0.5, 1.0, 2.0, and 10.0. *F*_obs_ was measured for each *f*_Co_ value over [Ni^II^]_total_ values between 0 μM and 250μM with [*P*]_total_ = 0.4 μM. We then generated binding curve predictions from the single-site model (Sect. 2.2) across the range of titrations for the *K*_D(Ni^II^)_, *K*_D(Co^II^)_, and *F*_max_ values determined in Sect. 3.1 (Table 1). The value of *F*_max_ used to parameterize the model was the average of the inferred values for Ni^II^ and Co^II^, which agree within one standard deviation, consistent with the hypothesis that they bind the same site. Across all *f*_Co_, we found the predicted binding curves adequately predicted the experimental data (Fig. 3). At larger *f*_Co_ values the experimental data reached saturation faster as a function of [Ni^II^]_total_ than at smaller *f*_Co_, which can be explained by the larger amount of total metal (Ni^II^ and Co^II^) for a given [Ni^II^]_total_ value and is consistent with the behaviour predicted by the model.

**Figure 3:**
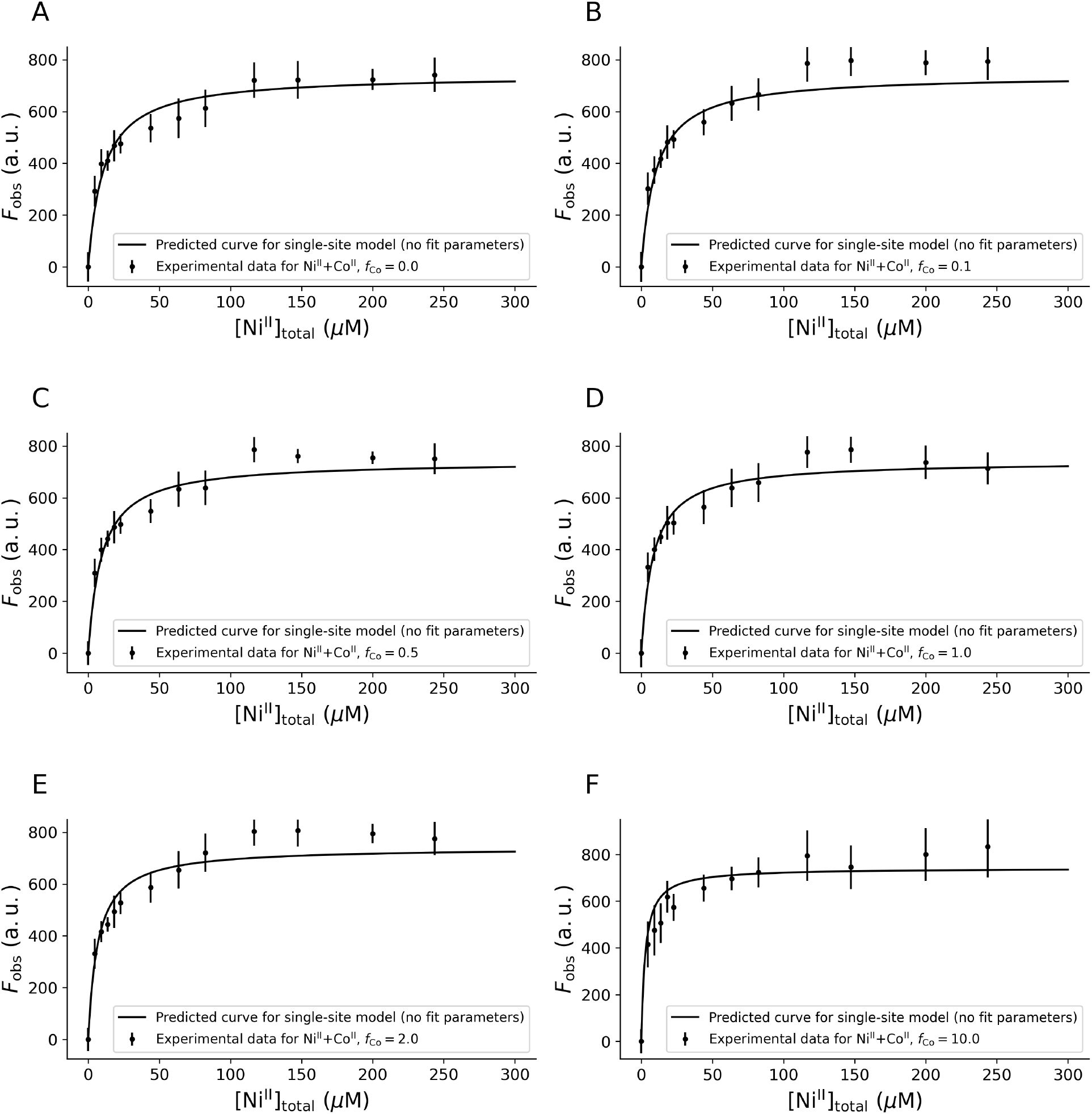
Model validation by prediction of binding curves for varying *f*_Co_. *Cc*NikZ-II was titrated with Ni^II^+Co^II^ mixtures of varying ratios (*f*_Co_ = 0.1 to 10) until saturation. The theoretically predicted (no fit) binding curves for the single-site model are plotted to evaluate the agreement of the model with the data. Experiments were performed in triplicate. The dots and error bars denote the mean and standard deviation, respectively, across the technical replicates for each titration step.

The single-site model formulated in (Sect. 2.2) does not account for different *F*_max_ values observed for *Cc*NikZ-II binding Ni^II^ compared to Zn^II^ (Table 1). Based on studies of the homologous protein *Cj*NikZ from *C. jejuni* [36], we expect *Cc*NikZ-II binds Ni^II^ with octahedral coordination geometry. In our previous study [32], we speculated that the different *F*_max_ exhibited by *Cc*NikZ-II when binding Ni^II^ compared to Zn^II^ may be due to Zn^II^ binding requiring a conformational change to accommodate its preference for forming tetrahedral metalloprotein complexes [6, 7], which may affect the polarity of the environment surrounding the proteinaceous tryptophans and, thereby, change the degree of fluoresence quenching upon binding. Modifying the single-site model to allow Ni^II^ and Zn^II^ binding events to result in different amounts of fluorescence quenching, however, still did not predict the Ni^II^+Zn^II^ data adequately (Supplementary Fig. S2 and Supplementary Table S4). Alternatively, the drastically different *F*_max_ values inferred for Zn^II^ compared to Ni^II^ (Table 1) may be suggestive of the existence of a second binding site on *Cc*NikZ-II with affinity for Zn^II^. This hypothesis is also consistent with Zn^II^’s preference for a tetrahedral geometry. Thus, we then assessed the ability of the two-site model (Sect. 2.3) to explain binding curves for *Cc*NikZ-II in solutions with Ni^II^ and Zn^II^ (Fig. 4).

**Figure 4:**
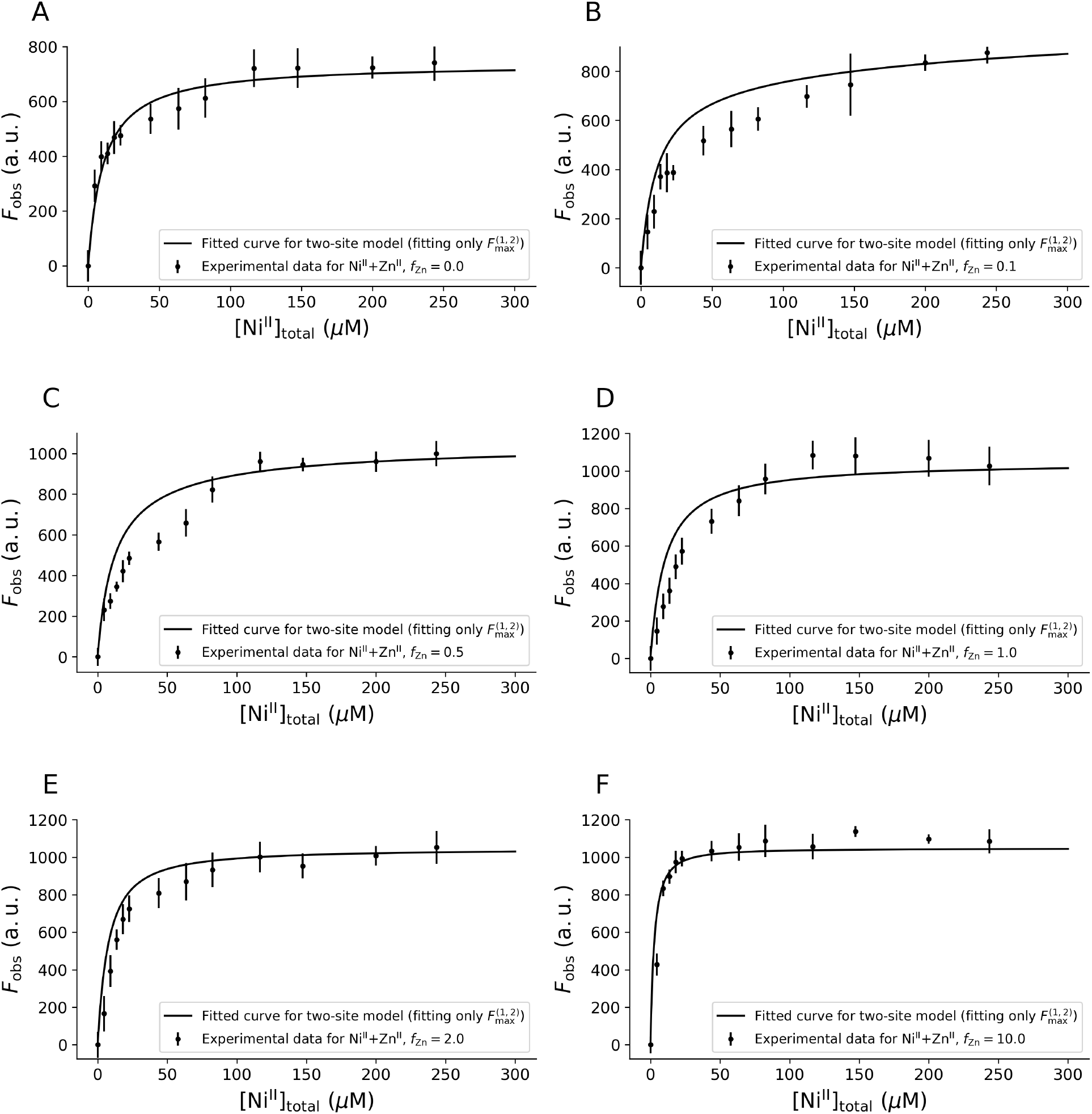
Model validation by prediction of binding curves for varying *f*_Zn_. *Cc*NikZ-II was titrated with Ni^II^+Zn^II^ mixtures of varying ratios (*f*_Zn_ = 0 to 10) until saturation. The fitted binding curves for the single-site model are plotted to evaluate the agreement of the model with the data. Note that the only free fit parameter is 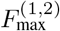, which was fitted to the data aggregated across all *f*_Zn_ values. Experiments were performed in triplicate. The dots and error bars denote the mean and standard deviation, respectively, across the technical replicates for each titration step.

To obtain binding curves for *Cc*NikZ-II in solutions with Ni^II^ and Zn^II^, we titrated the protein with metal stock solutions containing a mixture of the two metals at a fixed total metal ratios of *f*_Zn_ := [Zn^II^]_total_/[Ni^II^]_total_ = 0.1, 0.5, 1.0, 2.0, and 10.0. *F*_obs_ was measured for each *f*_Zn_ value over [Ni^II^]_total_ values between 0 μM and 250μM with [*P*]_total_ = 0.4 μM. We imposed the *K*_D(Ni^II^)_, *K*_D(Zn^II^)_, 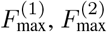 values determined in Sect. 3.1 (Table 1), leaving 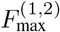 as the only free parameter in the two-site model for this system. Since 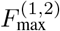 is the only free parameter and it should be an intrinsic property of the protein (*i.e*. independent of *f*_Zn_), we fit the two-site model to the data aggregated across all *f*_Zn_ values. Across all *f*_Zn_ values, we found adequate agreement between the resulting curves and the experimental data. Like in the Ni^II^+Co^II^ systems, at larger *f*_Zn_ values the experimental data reached saturation faster than at smaller *f*_Zn_, which can be explained by the larger amount of total metal (Ni^II^ and Zn^II^) for a given [Ni^II^]_total_ value and is consistent with the behaviour predicted by the model. However, unlike the Ni^II^+Co^II^ system, the experimental binding curves do not saturate to a constant value across all total metal ratios, a feature which was also captured by the fitted curves.

While we cannot rule out all possible alternative models to describe the data, we found that the single-site model (Sect. 2.2) adequately predicted the Ni^II^+Co^II^ data using parameter values obtained in independent experiments without any additional curve-fitting (Fig. 3). This provides further evidence of the validity of the inferred parameters from the single-metal experiments (Sect. 3.1) and suggests that the single-site binding model may be representative of the mechanistic details of the binding kinetics of *Cc*NikZ-II in the Ni^II^+Co^II^ system. Given the single-site model made adequate predictions for the Ni^II^+Co^II^ system, we did not evaluate the two-site model against the Ni^II^+Co^II^ data since the introduction of an additional fit parameter would trivially give rise to equal or better agreement with the data. Conversely, we found the single-site model did not adequately predict the Ni^II^+Zn^II^ data even when the difference in *F*_max_ values for each metal was accounted for (Supplementary Fig. S2 and Supplementary Table S4). However, we found the two-site (Sect. 2.2) model adequately explained the data (Fig. 4). While comparison between the two-site model and the Ni^II^+Zn^II^ data relied on fitting for 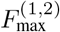, the adequate agreement of the fitted curves across each *f*_Zn_ value suggests the two-site model may describe the binding kinetics of *Cc*NikZ-II in the Ni^II^+Zn^II^ system. Given the adequate agreement of the proposed models and the data for both the Ni^II^+Co^II^ and Ni^II^+Zn^II^ systems, we proceeded with using the models to understand how changes in *K*_D(Ni^II^)_ and *K*_D(Co^II^)_ or *K*_D(Zn^II^)_ could alter the fraction of competitor-bound *Cc*NikZ-II in each system at thermodynamic equilibrium (Sect. 3.3), which could be used to guide protein engineering efforts towards more selective binders for target metals (in this case Ni^II^).

### 3.3 Model-based prediction of binding specificity over varying binding affinities

A goal of metalloprotein engineering is to improve a protein’s specificity towards a particular target metal. This objective can be met either by 1) increasing the protein’s binding affinity for the target metal (here, this corresponds to decreasing *K*_D(Ni^II^)_), or 2) decreasing its binding affinity for non-target metals (here, this corresponds to increasing *K*_D(Co^II^)_ or *K*_D(Zn^II^)_). While the qualitative effects of altering these affinities may be intuitive (increased affinity for target metal or decreased affinity for competitor metals reduces binding specificity), the degree to which the equilibrium fraction of competitor-bound protein complexes (here, given by [Co^II^]_bound_/[*P*]_total_ or [Zn^II^]_bound_/[*P*]_total_, where [Co^II^]_bound_ = [Co-*P*] and [Zn^II^]_bound_ = [Zn-*P*] + [Zn-*P*-Ni]) changes quantitatively with respect to changing *K*_D_ values is non-trivial.

For the Ni^II^+Co^II^ system, where the validated model (Sect. 3.2) predicts Ni^II^ and Co^II^ compete for a shared binding site on *Cc*NikZ-II (Sect. 2.2), we first considered the hypothetical case where *K*_D(Ni^II^)_ and *K*_D(Co^II^)_ can be altered independently. We used the the validated single-site binding model for the Ni^II^+Co^II^ system to generated theoretical predictions for the how the competition fraction [Co^II^]_bound_ changes as a function of the fold change in 1) *K*_D(Ni^II^)_ with *K*_D(Co^II^)_ kept constant at the value determined in Sect. 3.1 (Table 1) (Fig. 5A, C, E), and 2) *K*_D(Co^II^)_ with *K*_D(Ni^II^)_ kept constant at the value determined in Sect. 3.1 (Table 1) (Fig. 5B, D, F). We analyzed these predictions for [*P*]_total_ = 0.4 μM and *f*_Co_ = 0.1, 1, and 10 with [Ni]_total_ = 1 μM, 10 μM, 100 μM and in the limit [Ni]_total_ → ∞ μM (Fig. 5). The model predicted that a given fold change in *K*_D(Ni^II^)_ would give rise to comparable changes in the specificity fraction for an equivalent fold change in *K*_D(Co^II^)_, with the exception of sufficiently small [Ni^II^]_total_ values where the specificity fraction becomes marginally more sensitive to fold changes in *K*_D(Co^II^)_ (Fig. 5).

**Figure 5:**
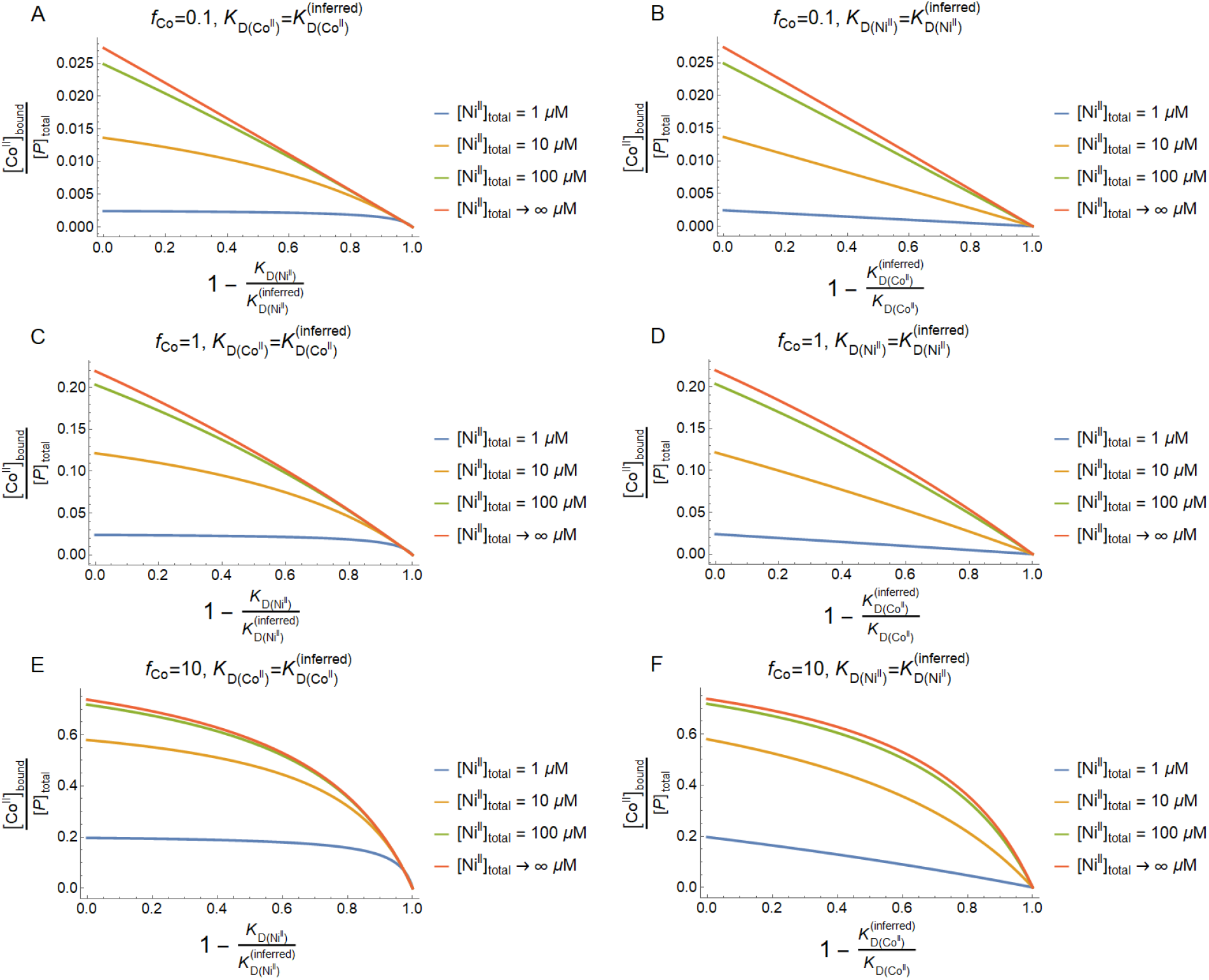
Predicting specificity fraction in Ni^II^+Co^II^ systems for individually varied metal binding affinities. Keeping *K*_D(Co^II^)_ constant at the value inferred via least squares optimization 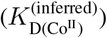 (Table 1), the specificity fraction is plotted as a function of *K*_D(Ni^II^)_ for [Ni^II^]_total_ = 1, 10, 100 μM across *f*_Co_ = 0.1, 1, 10 (A, C, E) for [*P*]_total_ = 0.4 μM. Similarly, keeping *K*_D(Ni^II^)_ constant at the value inferred via least squares optimization 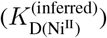 (Table 1), the specificity fraction is plotted as a function of *K*_D(Co^II^)_ for [Ni^II^]_total_ = 1, 10, 100 μM across *f*_Co_ = 0.1, 1, 10 (B, D, F) for [*P*]_total_ = 0.4 μM. The [Ni^II^]_total_ ≫ [*P*]_total_ limit of the specificity fraction specified by Eq. (17) is also shown for each *f*_Co_ value.

In practice it may be difficult to change *K*_D(Ni^II^)_ and *K*_D(Co^II^)_ independently when Ni^II^ and Co^II^ share the same binding site. Thus, we then considered how the specificity fraction changes when *K*_D(Ni^II^)_ and *K*_D(Co^II^)_ change together while holding their ratio constant. The ratio of dissociation constants for metals *M*_1_ and *M*_2_ defines the exchange constant *K*_ex_ for those metals and is commonly used to quantify the competition between *M*_1_ and *M*_2_ for binding the protein [17–19]. Taking *M*_1_ = Ni^II^ and *M*_2_ = Co^II^, the exchange constant is mathematically given as

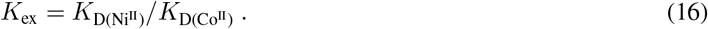

Intuitively, it is expected that a smaller *K*_ex_ corresponds to increased specificity for Ni^II^ relative to Co^II^. In the limit that [Ni^II^]_total_ ≫ [*P*]_total_ there is a simple quantitative relationship between the specificity fraction and *K*_ex_ for the single-site model (Sect. 2.2):

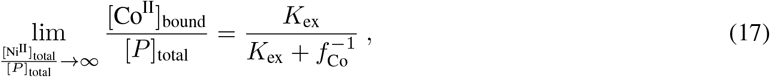

which confirms this intuition (see Supplementary Information Sect. S1.3 for derivation). The value of 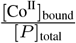 predicted by the single-site model for intermediate [Ni^II^]_total_ values also confirms this intuition (Fig. 5). However, for intermediate [Ni^II^]_total_ values, *K*_ex_ alone does not determine the specificity fraction for a fixed *f*_Co_ (Fig. 6). Moreover, this analysis of the single-site model predicts that the dissociation constant for the target metal (Ni^II^) should be increased to achieve improved specificity over the competitor (Co^II^) if their exchange constant remains constant.

**Figure 6:**
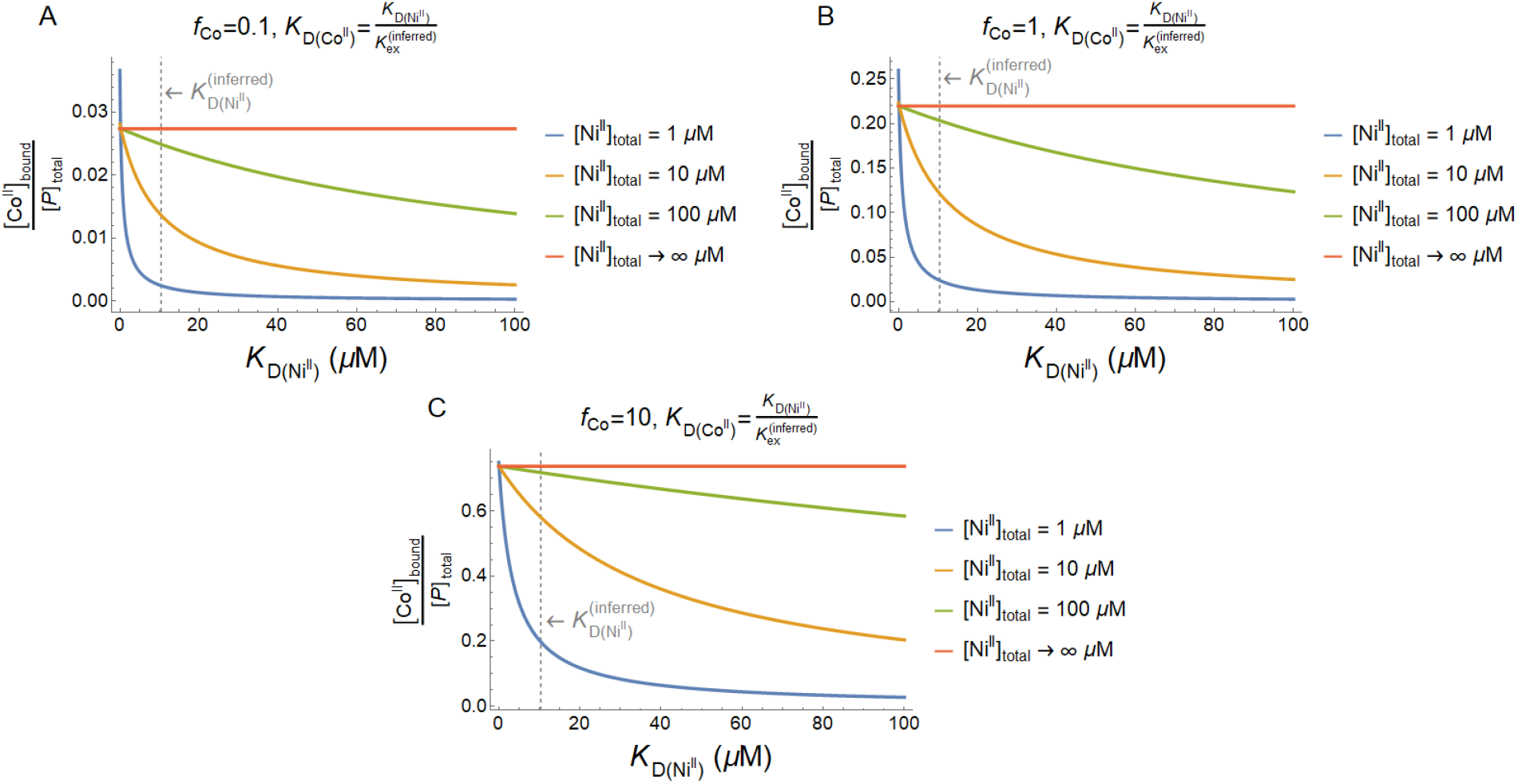
Predicting specificity fraction in Ni^II^+Co^II^ systems for varied metal binding affinities while fixing the exchange constant. Fixing *K*_ex_ at at the value calculated from *K*_*D*_ values inferred via least squares optimization 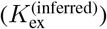 (Table 1), the specificity fraction is plotted as a function of *K*_D(Ni^II^)_ for [Ni^II^]_total_ = 1, 10, 100 μM across *f*_Co_ = 0.1, 1, 10 (A, B, C) for [*P*]_total_ = 0.4 μM. The [Ni^II^]_total_ ≫ [*P*]_total_ limit of the specificity fraction specified by Eq. (17) is also shown for each *f*_Co_ value.

For the Ni^II^+Zn^II^ system, where the validated model (Sect. 3.2) predicts Ni^II^ and Zn^II^ bind unique binding sites on *Cc*NikZ-II (Sect. 2.3), the model predicts changing the target metal’s dissociation constant (*K*_D(Ni^II^)_) has no effect on the fraction of Zn^II^-bound *Cc*NikZ-II complexes, while this quantity is decreased by increasing *K*_D(Zn^II^)_, as expected. In particular, since the binding of Ni^II^ and Zn^II^ are decoupled in such a model, both the fraction of Ni^II^-bound and Zn^II^-bound *Cc*NikZ-II is simply given by the bound-fraction expression Eq. (5) for the single-metal model (Sect. 3.1).

### 3.4 Limitations of the study

A protein’s *F*_max_ can differ between different metals due to differences in binding characteristics (*e.g*., differences in coordination geometry or binding sites). The *F*_max_ can also be affected by the protein’s environmental conditions (*e.g*., pH or temperature) and age. Earlier trials for the Ni^II^+Co^II^ experiments (not reported) resulted in binding curves that could not be adequately predicted by the specificity model Eq. (8), but was corrected by including a temperature equilibration step at 25°C prior to metal titrations. Additionally, *Cc*NikZ-II samples stored for long periods of time at 4 °C also experienced a reduction in the *F*_max_ (Supplementary Fig. S2), likely due to natural denaturation over time, similar to the activity of enzymes. This was corrected for by preparing fresh *Cc*NikZ-II solutions from freezer stocks. Failing to account for these conditions may result in different *F*_max_ values that may be incorrectly attributed to the nature of the binding event between a metal and protein.

There are also limitations related to model validation. While the binding curves predicted for the Ni^II^+Co^II^ experiments using the parameters inferred from the single-metal experiments appeared to predict the data adequately, the 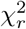 value for each single-metal fit was less than one (Table 1). This suggests either over-fitting or overestimated variance. We suggest the latter is likely due to each of these fits involving only two free parameters and the calculated 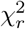 being less than one for the theoretical predictions (no fit) for most of the Ni^II^+Co^II^ experiments (Supplementary Table S1). We found that accounting for the different *F*_max_ values observed for Ni^II^ binding compared to Zn^II^ binding with a two-site binding model (Sect. 2.3) could reasonably explain the data (Fig. 4). However, the 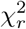 value for the two-site model fit to the Ni^II^+Zn^II^ data was greater than one for the aggregated data that 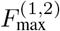 was estimated from (Table 2). This suggests either the fit did not fully capture the data or the data variance was underestimated. Likewise, the calculated 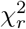 was greater than one when calculated for each *f*_Zn_ ≠ 0 (Supplementary Table S2). Given the previous arguments suggesting the data variance may be over-estimated, additional model features may be necessary to fully capture the data (*e.g*. cooperative binding resulting from allosteric interactions in the two-site model). Future work could explore these additional model complexities. Furthermore, while we showed reasonable agreement between our models and our data, it is impossible to rule out all possible alternative models. Future work could also explore additional validation approaches for the models, such as applying molecular dynamics simulations to verify stoichiometric and binding site multiplicity assumptions for particular multi-metal solutions.

## 4 Conclusion

In this work we extended the microplate-based intrinsic tryptophan fluorescence quenching ligand-titration assay [32] to study metalloprotein binding kinetics in solutions with multiple metals. Using this assay, we tested simple models describing the *in vitro* metal-binding kinetics of *Cc*NikZ-II in two-metal solutions. We found that different models were needed to adequately describe the measured binding curves in solutions consisting of Ni^II^+Co^II^ compared to Ni^II^+Zn^II^. Using these models, we quantitatively predicted how changes in binding affinity for each metal species affect the protein’s binding specificity for Ni^II^ in each two-metal solution and showed that the commonly used exchange constant does not uniquely determine binding specificity in general. This illustrates how mechanistic models could suggest design principles to guide the improvement of a metalloprotein’s binding specificity and that these principles depend on the composition of solutions in prospective bio-remediation applications. With emerging research on *de novo* proteins designed to have anti-Irving-William series properties [37, 38], our work here lays the groundwork for understanding how these synthetic proteins could be further engineered towards specific recovery of target metals from wastewater samples that contain other competing metals higher in the Irving-William series.

## CRediT authorship contribution statement

PD: Conceptualization, Experiments, Data curation, Methodology, Validation, Visualization, Writing - Original Draft, Review & Editing. BK: Conceptualization, Formal analysis, Methodology, Software, Validation, Visualization, Writing - Original Draft, Review & Editing. AY: Supervision, Funding acquisition, Writing - Review & Editing. AH: Supervision, Funding acquisition, Writing - Review & Editing. RM: Supervision, Funding acquisition, Writing - Review & Editing.

## Code & data availability

Data and code used for curve-fitting and calculating theoretical equilibrium concentrations is available on reasonable request from the corresponding author.

## Acknowledgments

We thank Shay Tal for providing constructive comments on an earlier version of the manuscript.

