## Supplementary Information for "Quantifying metal ion specificity of the nickel-binding protein *Cc*NikZ-II from *Clostridium carboxidivorans* in the presence of competing metal ions"

Patrick Diep[1], Brayden Kell[2,3], Alexander Yakunin[1,4], Andreas Hilfinger[2,3,5,6], and Radhakrishnan Mahadevan[1,7]

[1]BioZone Centre for Applied Bioscience and Bioengineering, Department of Chemical Engineering and Applied Chemistry, University of Toronto, Toronto, Canada
[2]Department of Physics, University of Toronto, Toronto, Canada
[3]Department of Chemical and Physical Sciences, University of Toronto Mississauga, Mississauga, Canada
[4]Centre for Environmental Biotechnology, Bangor University, Bangor, United Kingdom
[5]Department of Mathematics, University of Toronto, Toronto, Canada
[6]Department of Cell and Systems Biology, University of Toronto, Toronto, Canada
[7]Institute for Biomedical Engineering, University of Toronto, Toronto, Canada

In this supplement we provide mathematical derivations and additional figures and tables to complement the results in the Main Text.

**S2 Supplementary tables and figures** 4

### S1 Mathematical derivations

#### S1.1 Derivation of Main Text equation (8)

Here, we consider the single-site multi-metal model (Main Text Sect. 2.2) described by the set of coupled equilibrium reactions

$$M_iP \rightleftharpoons M_i + P \ , \qquad i = 1, \ \ldots, \ N \ . \tag{S1}$$

We wish to relate the fluorescence signal measured in an intrinsic tryptophan quenching assay for such a system to the equilibrium bound protein fraction. Previous work has given the following expression for the measured fluorescence $F$ for the special case of the above system with $N = 1$:

$$F = F_0(1 - \chi_b) + F_c\chi_b \ , \tag{S2}$$

where $F_0$ and $F_c$ are the fluorescence intensities measured when all protein molecules are free or bound, respectively, and $\chi_b$ is the fraction of protein molecules in a complex [1–3]. Under the assumption that each metal species has the same effect on the intrinsic tryptophan fluorescence upon binding, (S2) naturally holds for the general $N > 0$ case. Mathematically, this bound fraction is expressed

$$\chi_b = \frac{\sum_{i=1}^{N}[M_iP]}{[P]_{\text{total}}} \ . \tag{S3}$$

Simple algebraic manipulations of (S2) lead to

$$\frac{F_0 - F}{F_0 - F_c} = \chi_b \tag{S4}$$

Substituting (S3) leads to the desired expression:

$$\frac{F_{\text{obs}}}{F_{\text{max}}} = \frac{\sum_{i=1}^{N}[M_iP]}{[P]_{\text{total}}} \ , \tag{S5}$$

where we invoke the definitions $F_{\text{obs}} := F_0 - F$ and $F_{\text{max}} := F_0 - F_c$, as in the Main Text.

#### S1.2 Derivation of Main Text equation (15)

Here, we consider the two-site multi-metal model (Main Text Sect. 2.3) described by the set of coupled equilibrium reactions

$$\begin{aligned} M_iP &\rightleftharpoons M_i + P \ , \qquad i = 1, \ \dots, \ N \\ M_iM_jP &\rightleftharpoons M_i + M_j + P \ , \qquad (i,j) \in S_1 \times S_2 \ , \end{aligned} \tag{S6}$$

where $\times$ denotes the Cartesian product (*i.e.* $(i,j) \in S_1 \times S_2 \iff i \in S_1, \ j \in S_2$) and the sets $S_1$ and $S_2$ are defined as follows:

$$\begin{aligned} S_1 &:= \{i \mid M_i \text{ binds site } 1\} \\ S_2 &:= \{i \mid M_i \text{ binds site } 2\} \end{aligned} \tag{S7}$$

We assume that each species that binds site 1 has an equal quenching effect on the protein's intrinsic tryptophan fluorescence and likewise for site 2, but that the effects across the two sites may differ. Moreover, we allow for the formation of the a second-order complex $M_iM_jP$ to have an overall effect on the protein's intrinsic tryptophan fluorescence that may not necessarily equal the sum of the quenching effects from each single metal binding its corresponding site on the protein in its fully unbound state. Under these assumptions, (S2) generalizes to

$$F = F_0(1 - (\chi_b^{(1)} + \chi_b^{(2)} + \chi_b^{(1,2)})) + F_c^{(1)}\chi_b^{(1)} + F_c^{(2)}\chi_b^{(2)} + F_c^{(1,2)}\chi_b^{(1,2)}, \tag{S8}$$

where $\chi_b^{(1)}$, $\chi_b^{(2)}$ correspond to the fraction of protein molecules with only site 1 occupied or only site 2 occupied, respectively, and $\chi_b^{(1,2)}$ corresponds to the fraction of the protein molecules with both sites occupied. $F_c^{(1)}$, $F_c^{(2)}$ represent the fluorescence when all protein molecules have only site 1 occupied or only site 2 occupied, respectively, and $F_c^{(1,2)}$ represents the fluorescence when all protein molecules have both sites occupied. Mathematically, the fractions of different complex types (only site 1 occupied, only site 2 occupied, and both sites occupied) are expressed

$$\begin{aligned} \chi_b^{(1)} &= \frac{\sum_{i \in S_1}[M_iP]}{[P]_{\text{total}}} \\ \chi_b^{(2)} &= \frac{\sum_{i \in S_2}[M_iP]}{[P]_{\text{total}}} \\ \chi_b^{(1,2)} &= \frac{\sum_{(i,j) \in S_1 \times S_2}[M_iM_jP]}{[P]_{\text{total}}} \ . \end{aligned} \tag{S9}$$

Simple algebraic manipulations of (S8) lead to

$$\underbrace{F_0 - F}_{F_{\text{obs}}} = \underbrace{(F_0 - F_c^{(1)})}_{=:F_{\text{max}}^{(1)}}\chi_b^{(1)} + \underbrace{(F_0 - F_c^{(2)})}_{=:F_{\text{max}}^{(2)}}\chi_b^{(2)} + \underbrace{(F_0 - F_c^{(1,2)})}_{=:F_{\text{max}}^{(1,2)}}\chi_b^{(1,2)} \ . \tag{S10}$$

Substituting (S9) leads to the desired expression:

$$F_{\text{obs}} = \frac{F_{\text{max}}^{(1)}\sum_{i \in S_1}[M_iP] + F_{\text{max}}^{(2)}\sum_{i \in S_2}[M_iP] + F_{\text{max}}^{(1,2)}\sum_{(i,j) \in S_1 \times S_2}[M_iM_jP]}{[P]_{\text{total}}} \ . \tag{S11}$$

#### S1.3 Derivation of Main Text equation (17)

Here, we consider the single-site multi-metal model (Main Text Sect. 2.2) described by the set of coupled equilibrium reactions

$$M_iP \rightleftharpoons M_i + P \ , \qquad i = 1, \ \dots, \ N \ . \tag{S12}$$

Without loss of generality, let $M_1$ denote the metal of interest such that the competitor-bound protein fraction is given as $[M_{i \neq 1}]_{\text{bound}}/[P]_{\text{total}} = \sum_{i=2}^{N}[M_iP]/[P]_{\text{total}}$. We are interested in calculating this quantity in the limit that

$[M_1]_{\text{total}} \gg [P]_{\text{total}}$ with the total metal ratios $f_i \coloneqq [M_i]_{\text{total}}/[M_1]_{\text{total}}$ held constant. In this limit nearly all protein will be bound by a metal, leading to the mass balance approximations

$$[P]_{\text{total}} \approx \sum_{i=1}^{N}[M_iP] \tag{S13}$$

$$[M_i]_{\text{total}} \approx [M_i] \ . \tag{S14}$$

From the definition of the dissociation constant we then have

$$\frac{K_{\text{D}(i)}}{K_{\text{D}(1)}} = \frac{[M_i][M_1P]}{[M_1][M_iP]} \tag{S15}$$

$$\approx f_i \frac{[M_1P]}{[M_iP]} \ . \tag{S16}$$

Isolating for $[M_iP]$ leads and summing over all $i > 1$ then leads to

$$\sum_{i=2}^{N}[M_iP] \approx K_{\text{D}(1)}[M_1P]\sum_{i=2}^{N}\frac{f_i}{K_{\text{D}(i)}} \ . \tag{S17}$$

From (S14), we have $[M_1P] \approx [P]_{\text{total}} - \sum_{i=2}^{N}[M_iP]$. Substituting this the above expression then and solving for $\sum_{i=2}^{N}[M_iP]$ gives

$$\sum_{i=2}^{N}[M_iP] \approx [P]_{\text{total}}\left(1 + K_{\text{D}(1)}\sum_{i=2}^{N}\frac{f_i}{K_{\text{D}(i)}}\right)^{-1} K_{\text{D}(1)}\sum_{i=2}^{N}\frac{f_i}{K_{\text{D}(i)}} \ . \tag{S18}$$

Equivalently, we have

$$\frac{[M_{i\neq 1}]_{\text{bound}}}{[P]_{\text{total}}} \approx \frac{K_{\text{D}(1)}}{K_{\text{D}(1)} + \left(\sum_{i=2}^{N}\frac{f_i}{K_{\text{D}(i)}}\right)^{-1}} \ , \tag{S19}$$

which approximates the competitor-bound protein fraction in the limit of total metal concentrations much larger than the total protein concentration and is exact in the formal limit $[M_1]_{\text{total}}/[P]_{\text{total}} \to \infty$ for $f_i > 0$ constant. This is the general form of equation (17) from the Main Text for arbitrary metals of interest and arbitrarily many competing metals. To recover equation (17) from the Main Text exactly, one takes $M_1 = \text{Ni}^{\text{II}}$, $M_1 = \text{Co}^{\text{II}}$ and invokes the exchange coefficient definition $K_{\text{ex}} = K_{\text{D}(\text{Ni}^{\text{II}})}/K_{\text{D}(\text{Co}^{\text{II}})}$.

#### S1.4 Derivation of theoretical equilibrium concentrations

##### S1.4.1 Single-site binding model for multi-metal solutions

Here, we consider the single-site multi-metal model (Main Text Sect. 2.2) described by the set of coupled equilibrium reactions

$$M_iP \rightleftharpoons M_i + P \ , \qquad i = 1,\ \dots,\ N \ . \tag{S20}$$

Denoting the total concentration of protein and each metal species as $[P]_{\text{total}}$ and $[M_i]_{\text{total}}$, respectively, the mass balance equations for such a system are given as

$$\begin{aligned} [P]_{\text{total}} &= [P] + \sum_{i=1}^{N}[M_iP] \\ [M_i]_{\text{total}} &= [M_i] + [M_iP] \ , \qquad i = 1,\ \dots,\ N \ , \end{aligned} \tag{S21}$$

where $[\cdot]$ denotes an equilibrium concentration. Additionally, the dissociation constants for each species are defined as

$$K_{\text{D}(i)} = \frac{[M_i][P]}{[M_iP]} \ , \qquad i = 1,\ \dots,\ N \ . \tag{S22}$$

Treating the $K_{\text{D}(i)}$ values as (known) model parameters, the system of equations (S21) with (S22) is algebraically closed such that the equilibrium concentrations $[P]$, $[M_i]$ ($i = 1,\ \dots,\ N$) may be obtained. To obtain theoretical predictions for the $\text{Ni}^{\text{II}}$+$\text{Co}^{\text{II}}$ system, we used the `Solve[]` function in `Wolfram Mathematica 12` to obtain closed-form parametric solutions to this set of equations for the case of $N = 2$ and then substituted the relevant values for the total concentrations and dissociation constants (see Main Text Sect. 2.4). In general, for an arbitrary $N \in \mathbb{Z}_{>0}$, solving the system of equations (S21) with (S22) for arbitrary values of the total concentrations and dissociation constants corresponds to solving for the roots of a $N + 1$-degree polynomial with general coefficients. Thus, by the Abel–Ruffini theorem, closed-form parametric solutions for the equilibrium concentrations may be obtained only for $N \leq 3$ [4]. For researchers interested in systems with $N > 3$, numerical solutions may be obtained for particular values of the total concentrations and dissociation constants [5, 6].

##### S1.4.2 Two-site binding model for multi-metal solutions

Here, we consider the two-site multi-metal model (Main Text Sect. 2.3) described by the set of coupled equilibrium reactions

$$\begin{aligned} M_iP &\rightleftharpoons M_i + P \ , \qquad i = 1, \ \ldots, \ N \\ M_iM_jP &\rightleftharpoons M_i + M_j + P \ , \qquad (i,j) \in S_1 \times S_2 \ , \end{aligned} \tag{S23}$$

where $\times$ denotes the Cartesian product (*i.e.* $(i,j) \in S_1 \times S_2 \iff i \in S_1, \ j \in S_2$) and the sets $S_1$ and $S_2$ are defined as follows:

$$\begin{aligned} S_1 &:= \{i \mid M_i \text{ binds site 1}\} \\ S_2 &:= \{i \mid M_i \text{ binds site 2}\} \end{aligned} \tag{S24}$$

Denoting the total concentration of protein and each metal species as $[P]_{\text{total}}$ and $[M_i]_{\text{total}}$, respectively, the mass balance equations for such a system are given as

$$\begin{aligned} [P]_{\text{total}} &= [P] + \sum_{i=1}^{N} [M_iP] + \sum_{(i,j)\in S_1 \times S_2} [M_iM_jP] \\ [M_i]_{\text{total}} &= [M_i] + [M_iP] \ , \qquad i = 1, \ \ldots, \ N \ . \end{aligned} \tag{S25}$$

Additionally, the dissociation constants for each species are defined as

$$K_{\text{D}(i)} = \frac{[M_i][P]}{[M_iP]} \ , \qquad i = 1, \ \ldots, \ N \ . \tag{S26}$$

We assume an absence of allostery, such that the equilibrium constants for the reactions $M_iM_jP \rightleftharpoons M_i + M_j + P$ are constrained by

$$\frac{[M_i][M_j][P]}{[M_iM_jP]} = K_{\text{D}(i)} K_{\text{D}(j)} \ , \qquad (i,j) \in S_1 \times S_2 \ . \tag{S27}$$

Treating the $K_{\text{D}(i)}$ values as (known) model parameters, the system of equations (S25) with (S26) and (S27) is algebraically closed such that the equilibrium concentrations $[P]$, $[M_i]$ $(i = 1, \ \ldots, \ N)$ may be obtained. To obtain theoretical predictions for the $Ni^{II}$+$Zn^{II}$ system, we used the `Solve[]` function in `Wolfram Mathematica 12` to obtain closed-form parametric solutions to this set of equations for the case of $N = 2$ and then substituted the relevant values for the total concentrations and dissociation constants (see Main Text Sect. 2.4). Like the single-site model, closed-form parametric solutions for the equilibrium concentrations may be obtained only for $N \leq 3$ but the numerical approach [6] may still be applied to obtain numerical solutions to this model for $N > 3$.

### S2 Supplementary tables and figures

Supplementary Table S1: $\chi^2_r$ for Single-site Model Predictions for $Ni^{II}$+$Co^{II}$ Experiments

| | $f_{\text{Co}}$ | $\chi^2_r$ |
|---|---|---|
| $Ni^{II}$+$Co^{II}$ experiments | 0 | 0.5 |
| (Main Text Fig. 3) | 0.1 | 0.9 |
| | 0.5 | 1.4 |
| | 1 | 0.6 |
| | 2 | 1.4 |
| | 10 | 0.6 |

Supplementary Table S2: $\chi^2_r$ for Two-site Model Fits for $Ni^{II}$+$Zn^{II}$ Experiments

| | $f_{\text{Zn}}$ | $\chi^2_r$ |
|---|---|---|
| $Ni^{II}$+$Zn^{II}$ experiments | 0 | 0.5 |
| (Main Text Fig. 4) | 0.1 | 4.2 |
| | 0.5 | 10.5 |
| | 1 | 3.9 |
| | 2 | 2.1 |
| | 10 | 3.3 |

Supplementary Table S3: Mean total recovery fractions for $Ni^{II}$ stoichiometry experiments

| | | Mean total $Ni^{II}$ recovery % | Mean total *Cc*NikZ-II recovery % |
|---|---|---|---|
| $Ni^{II}$ stoichiometry experiments (Main Text Fig. 1AB) | Negative control | 0.31 | × |
| | Only $Ni^{II}$ | 71.02 | × |
| | Only *Cc*NikZ-II | 0.54 | 121.64 |
| | $Ni^{II}$ & *Cc*NikZ-II | 80.00 | 109.07 |

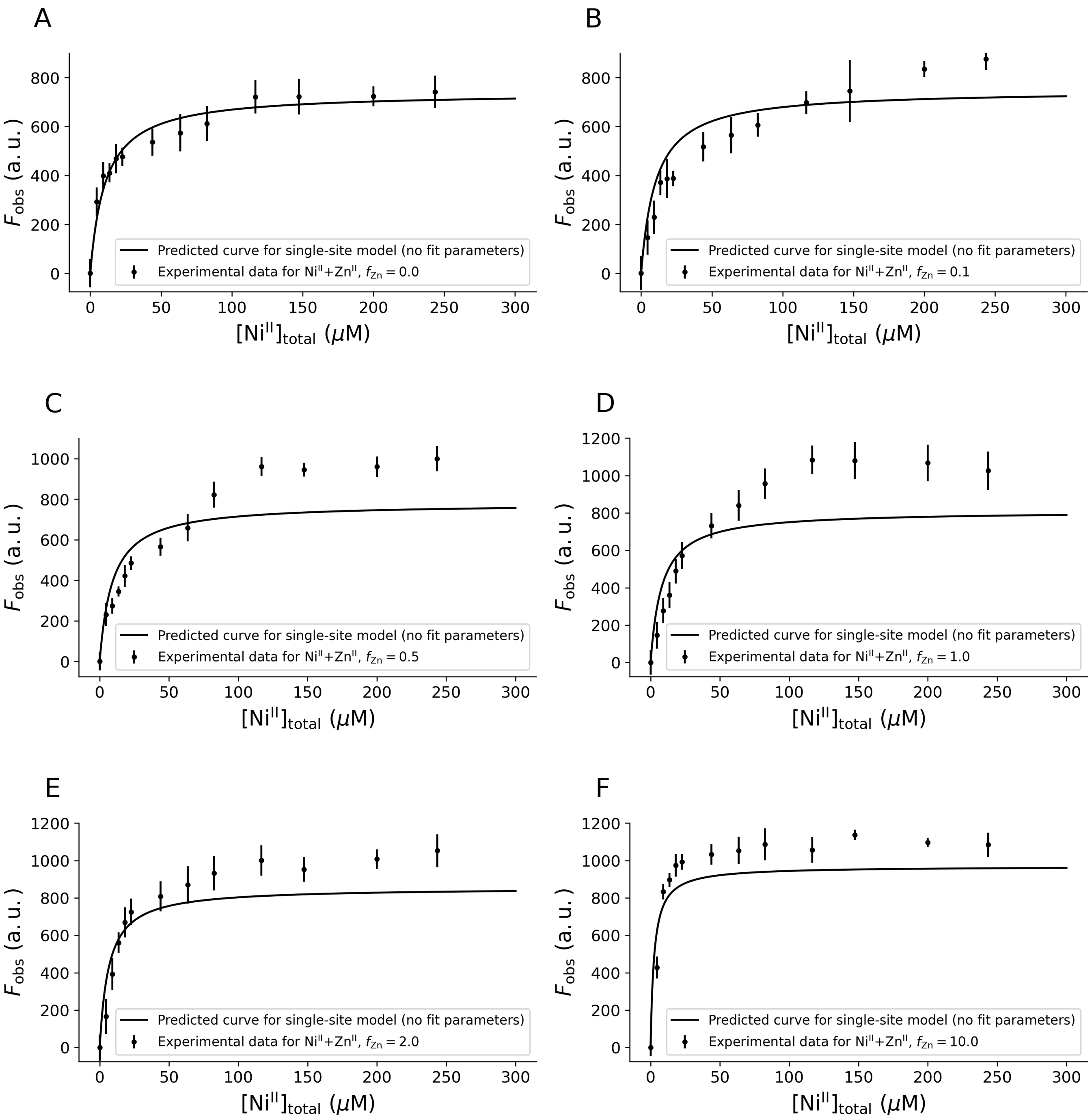

Supplementary Figure S1: **Model rejection by prediction of binding curves for varying $f_{Zn}$.** *Cc*NikZ-II was titrated with $Ni^{II}+Zn^{II}$ mixtures of varying ratios ($f_{Zn} = 0$ to 10) until saturation. The theoretically predicted binding curves for the single-site model are plotted to evaluate the agreement of the model with the data. Here, the single-site model was modified to account for the different $F_{max}$ values observed in the single-ligand experiments (Main Text Table 1) such that $F_{obs} = (F_{max}^{(Ni^{II})}[Ni^{II}\text{-}P] + F_{max}^{(Zn^{II})}[Zn^{II}\text{-}P])/[P]_{total}$, where $F_{max}^{(Ni^{II})}$ and $F_{max}^{(Zn^{II})}$ were the $F_{max}$ values obtained in the single-ligand experiments (Main Text Table 1). Experiments were performed in triplicate. The dots and error bars denote the mean and standard deviation, respectively, across the technical replicates for each titration step.

Supplementary Table S4: $\chi_r^2$ for Single-site Model Predictions for $Ni^{II}$+$Zn^{II}$ Experiments

| | $f_{Zn}$ | $\chi_r^2$ |
|---|---|---|
| $Ni^{II}$+$Zn^{II}$ experiments (Supplementary Fig. S1) | 0 | 0.5 |
| | 0.1 | 4.2 |
| | 0.5 | 11.2 |
| | 1 | 5.2 |
| | 2 | 2.9 |
| | 10 | 9.3 |

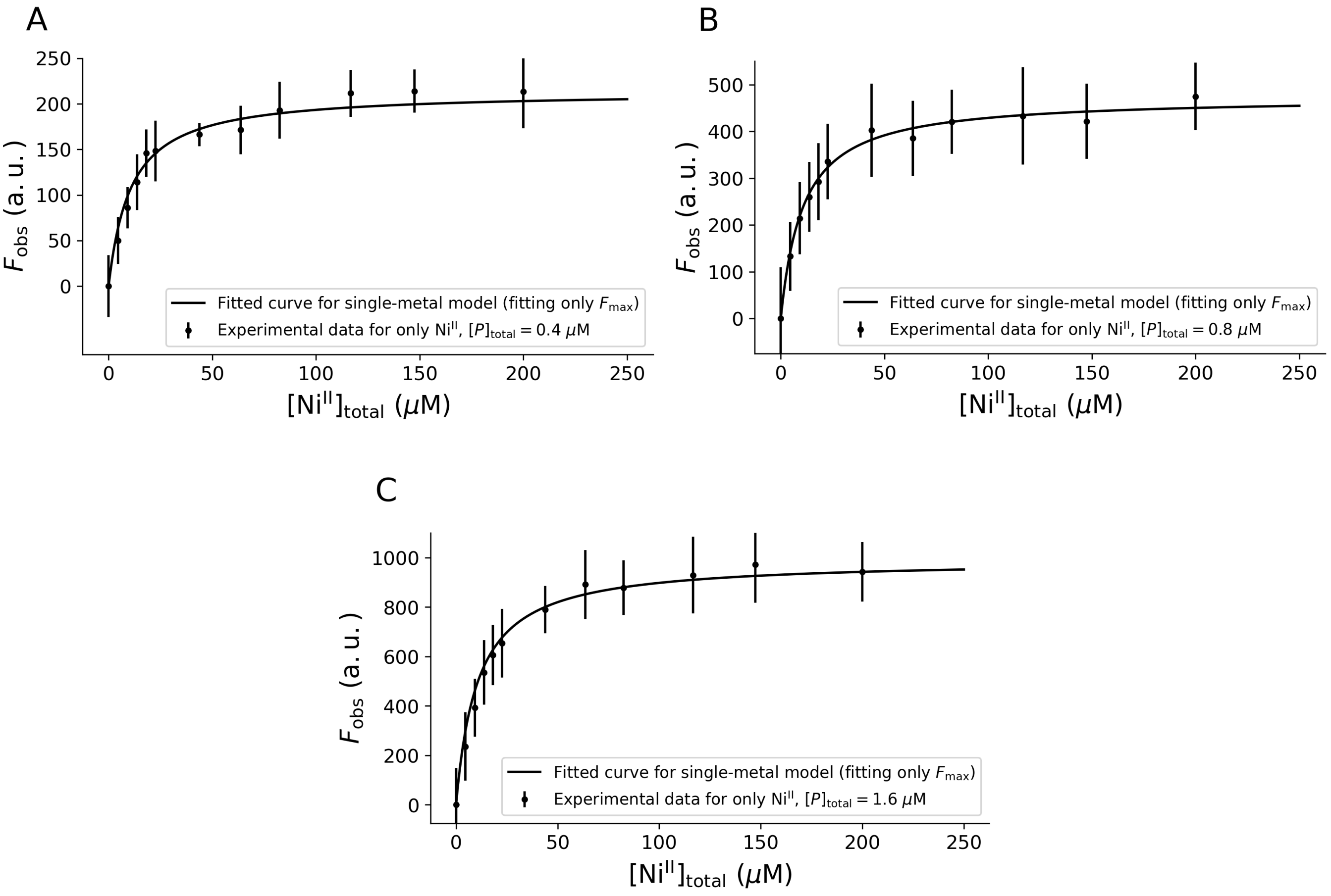

Supplementary Figure S2: **Fitted binding curves for varied $[P]_{total}$ values.** *Cc*NikZ-II was titrated with $Ni^{II}$ at 0.4 µM (A), 0.8 µM (B), and 1.6 µM (C). The binding curves were obtained by substituting the $K_D$ value obtained for $Ni^{II}$ from an independent experiment for $[P]_{total} = 0.4$ µM (Main Text Table 1) and fitting for $F_{max}$ in the single-metal metal model (Main Text Sect. 2.1) for each $[P]_{total}$ value. Experiments were performed in triplicate. The dots and error bars denote the mean and standard deviation, respectively, across the technical replicates for each titration step. We expect the smaller $F_{max}$ value for $[P]_{total} = 0.4$ µM observed here (Supplementary Table S5) compared to the value inferred in the Main Text (Main Text Table 1) is due to natural protein denaturation after being stored for a long period at 4°C (see Main Text Sect. 3.4).

Supplementary Table S5: Fitted Parameter and $\chi_r^2$ for $Ni^{II}$ Experiments with Varied $[P]_{total}$

| | $[P]_{total}$ | $F_{max}$ | $\chi_r^2$ |
|---|---|---|---|
| $Ni^{II}$ experiments (Fig. S2) | 0.4 µM | 213.7 ± 4.2 | 0.2 |
| | 0.8 µM | 473.7 ± 5.7 | 0.03 |
| | 1.6 µM | 991.7 ± 11.1 | 0.05 |
